# Accuracy of the pneumatic method for estimating xylem vulnerability to embolism in temperate diffuse-porous tree species

**DOI:** 10.1101/2021.02.15.431295

**Authors:** Sharath S. Paligi, Roman M. Link, Emilie Isasa, Paulo Bittencourt, Juliano Sarmento Cabral, Steven Jansen, Rafael S. Oliveira, Luciano Pereira, Bernhard Schuldt

## Abstract

- The increasing frequency of global change-type droughts has created a need for fast, accurate and widely applicable techniques for estimating xylem embolism resistance to improve forecasts of future forest changes.
- We used data from 12 diffuse-porous temperate tree species covering a wide range of xylem safety to compare the pneumatic and flow-centrifuge method for constructing xylem vulnerability curves. We evaluated the agreement between parameters estimated with both methods and the sensitivity of pneumatic measurements to the measurement duration.
- The agreement between xylem water potentials at 50% air discharged (PAD) estimated with the Pneumatron and 50% loss of hydraulic conductivity (PLC) estimated with the flow-centrifuge method was high (mean signed deviation: 0.12 MPa, Pearson correlation: 0.96 after 15 sec of gas extraction). However, the relation between the estimated slopes was more variable, resulting in lower agreement in xylem water potential at 12% and 88% PAD/PLC. All parameters were sensitive to the duration of the pneumatic measurement, with highest overall agreement between methods after 16 sec.
- We conclude that, if applied correctly, the pneumatic method enables fast and inexpensive estimations of embolism resistance for a wide range of temperate, diffuse-porous species, which makes it attractive for predicting plant performance under climate change.

## Introduction

In the last decades, unprecedented climate fluctuations and the resulting extreme drought events have led to large-scale tree dieback events worldwide (Allen *et al*., 2010, 2015; Brando *et al*., 2019). With the global rise in frequency, intensity and duration of drought spells predicted by current climate projections (cf. Field *et al*., 2012; Trenberth *et al*., 2014), large-scale drought-induced tree mortality events become increasingly likely (Brodribb *et al*., 2020).

To improve the prediction of demographic and compositional changes in forest ecosystems, a better understanding of physiological mechanisms associated with the death of trees in response to drought is necessary (Allen *et al*., 2010; McDowell *et al*., 2013a, 2013b). In this context, traits that quantify the vulnerability of a tree’s xylem to drought-induced embolism have received particular attention (Choat *et al*., 2018; Brodribb *et al*., 2020). Vulnerability to embolism is usually expressed by the parameters of xylem vulnerability curves (VCs), i.e. curves describing the consecutive loss of hydraulic conductance (percent loss of conductivity, PLC) as a function of increasingly negative xylem pressures (cf. Sperry *et al*., 1988; Cochard *et al*., 2013). Most commonly, VCs are described by the water potential at 12, 50 or 88% loss of hydraulic conductance (*P*_12_, *P*_50_ and *P*_88_, respectively) and the slope of the curve at one of the respective locations. The parameters of VCs have been linked to mechanistic thresholds for xylem functioning (cf. Brodribb & Cochard, 2009; Urli *et al*., 2013; Delzon & Cochard, 2014), and are closely coordinated with stomatal regulation (Martin-StPaul *et al*., 2017). Across biomes, xylem embolism resistance has been associated with the susceptibility of a species to drought-induced mortality (Anderegg *et al*., 2016; Adams *et al*., 2017; Correia *et al*., 2019; Powers *et al*., 2020) and thus mirrors the distribution of species along aridity gradients (Blackman *et al*., 2014; Trueba *et al*., 2017; Oliveira *et al*., 2019). Due to their immediate mechanistic interpretation, VC parameters and derived quantities such as hydraulic safety margins (Meinzer *et al*., 2009) are increasingly incorporated in process-based vegetation models to describe plant drought responses and associated drought-induced tree mortality (McDowell *et al*., 2013a, 2013b; Christoffersen *et al*., 2016; Xu *et al*., 2016; Davi & Cailleret 2017; Eller *et al*., 2020).

Xylem VCs can be established through a large number of different techniques, such as the bench dehydration (Sperry *et al*., 1988), air injection (Cochard *et al*., 1992), flow-centrifuge (Cochard *et al*., 2005), micro-CT (Brodersen *et al*., 2010), pneumatic (Pereira *et al*., 2016), optical (Brodribb *et al*., 2016) and relative water loss method (Rosner *et al*., 2019). However, none of these methods is unequivocally reliable for angiosperm and gymnosperm species, either due to measurement artefacts associated with vessel length and porosity (Cochard *et al*., 2013; Jansen *et al*., 2015) or because they are not suitable for rapid measurements of large number of samples (Cochard *et al*., 2013; Nolf *et al*., 2017). Given their usefulness to predictive models, methods for the measurement of xylem embolism that are simple, accessible, reliable and applicable for a wide range of taxonomic groups and xylem types are needed.

A novel, promising route for fast indirect VC measurements is the pneumatic method (Pereira *et al*., 2016; Jansen *et al*., 2020), which estimates the amount of xylem embolism by measuring the increase of air volume in the xylem in bench-dried plant samples with increasingly negative xylem pressure. Recently, Pereira *et al*. (2020) proposed an automated device, the Pneumatron, which automatically measures air discharge from a plant sample at a high temporal resolution and permits a high sample throughput. As the measurement principle of the Pneumatron does not directly depend on the measurement of xylem water transport, and embolism is induced by bench dehydration, it is assumed to be relatively robust against measurement artefacts related to sample excision and preparation as well as vessel length related artefacts (Pereira *et al*., 2016, 2020), which are known to affect several hydraulic VC methods (cf. Choat *et al*., 2010; Wheeler *et al*., 2013; Martin-StPaul *et al*., 2014; Torres-Ruiz *et al*., 2014, 2015).

The pneumatic method has already been applied to construct VCs for branches, leaves and roots of tropical, subtropical and temperate species, covering diffuse-porous, ring-porous and coniferous species (Pereira *et al*., 2016, 2020; Zhang *et al*., 2018; Wu *et al*., 2020; Sergent *et al*., 2020). The results from these studies indicate the suitability of the method for diffuse-porous (Pereira *et al*., 2016; Zhang *et al*., 2018; Sergent *et al*., 2020) and for ring-porous species (though the latter only after gluing growth rings other than the current year rings; cf. Zhang *et al*., 2018). However, Zhang *et al*. (2018) reported that the pneumatic method resulted in lower estimates of embolism resistance of two conifer species compared to the flow-centrifuge method. Similarly, in a recent methodological comparison of VC techniques, Sergent *et al*. (2020) found the pneumatic method to result in lower embolism resistance estimates compared to other methods in further two conifer species, and reported inconsistent estimates for long-vesseled species. Several sources of uncertainty have been identified for the pneumatic method that leave room for potential methodological improvements. Notably, its estimates are known to be sensitive to the choice of the reservoir volume and the duration of the air discharge measurement (Pereira *et al*., 2016, 2020). Prior studies have used vastly different time intervals to measure air discharge (AD) from the xylem segment, ranging from 150 sec (Pereira *et al*., 2016; Chen *et al*., 2021), over 120 sec (Zhang *et al*., 2018; Sergent *et al*., 2020; Wu *et al*., 2020), to only 30 sec (Pereira *et al*., 2020). Since a 15 sec duration was predicted as optimal discharge time based on the Unit Pipe Pneumatic model (Yang *et al*., 2021), there is a need to test this hypothesis experimentally. Moreover, despite considerable attention to measuring artefacts and methodological concerns, we currently lack a rigorous statistical framework to compare embolism resistance methods.

This study uses a dataset of vulnerability curve measurements from 36 trees belonging to 12 temperate, diffuse-porous tree species to i) assess how well the parameters of the vulnerability curves obtained with the pneumatic method (Pneumatron) agree with estimates obtained from the flow-centrifuge method (Cavitron) in terms of systematic deviations, random deviations and overall agreement, and to ii) identify the optimal duration for air discharge measurements.

## Materials and Methods

### Plant material

Plant material from trees belonging to 12 temperate diffuse-porous tree species (Table 1) was collected between mid-July and mid-September 2019 from a nursery in Veitshöchheim (49°50’24.3”N, 9°52’38.4”E) near Würzburg, Germany. These trees were planted in the year 2011 in the framework of a long-term comparative experiment designed to identify appropriate tree species for urban planting. The plant material cultivated in Veitshöchheim was obtained from selected nurseries across Central Europe. Due to anomalies in the data for some of the species (see below), additional branch samples were obtained from adult *Tilia cordata* and *Tilia platyphyllos* trees growing at Ulm University, Germany (48°25’20.3”N, 9°57’20.2”E) and of *Tilia japonica* from the Würzburg botanical garden (49°45’56.7”N 9°55’58.1”E) in September 2020.

**Table 1:** List of the 12 diffuse-porous tree species used in the present study, average midday leaf water potential (*ψ*_midday_) measured in August 2020 and the average diameter at breast height (DBH) of the selected trees per species (mean ± SE); *n*_P_ and *n*_H_ indicate the number of xylem vulnerability curves measured for the pneumatic and the flow-centrifuge method, respectively (values in brackets indicate when branch samples were collected from a single tree). Asterisks (*) indicate xylem pressures measured with stem psychrometers; hashes (#) indicate individuals from the same species measured with the pressure bomb.

| Species | Family | $n_P$ | $n_H$ | $\Psi_{\text{midday}}$ (MPa) | DBH (cm) |
| --- | --- | --- | --- | --- | --- |
| *Betula pendula* | Betulaceae | 3 | 3 | $-1.60 \pm 0.04$ | $10.93 \pm 0.64$ |
| *Betula utilis* | Betulaceae | 3 | 3 | $-1.54 \pm 0.06$ | $08.88 \pm 0.42$ |
| *Carpinus betulus* | Betulaceae | 3 | 3 | $-2.47 \pm 0.06$ | $10.75 \pm 0.44$ |
| *Crataegus persimilis* | Rosaceae | 3 | 3 | $-3.42 \pm 0.18$ | $07.55 \pm 0.16$ |
| *Ostrya carpinifolia* | Betulaceae | 3 | 3 | $-2.97 \pm 0.11$ | $09.18 \pm 0.11$ |
| *Platanus x acerifolia* | Platanaceae | 3 | 3 | $-1.74 \pm 0.03$ | $10.00 \pm 0.24$ |
| *Platanus orientalis* | Platanaceae | 3 | 3 | $-1.51 \pm 0.04$ | $12.07 \pm 0.35$ |
| *Pyrus calleryana* | Rosaceae | 2 | 3 | $-3.52 \pm 0.30$ | $11.88 \pm 0.32$ |
| *Sorbus latifolia* | Rosaceae | 3 | 3 | $-3.64 \pm 0.19$ | $09.30 \pm 0.39$ |
| *Tilia cordata* * | Malvaceae | 5(1) | 5(1) | NA | $28.00 \pm 0.00$ |
| *Tilia cordata*# | Malvaceae | 3 | 3 | $-1.90 \pm 0.04$ | $11.40 \pm 0.46$ |
| *Tilia japonica* | Malvaceae | 3(1) | 4(1) | NA | $30.40 \pm 0.00$ |
| *Tilia platyphyllos* * | Malvaceae | 4(1) | 7(1) | NA | $70.00 \pm 0.21$ |
| *Tilia platyphyllos*# | Malvaceae | 3 | 3 | $-1.80 \pm 0.08$ | $12.38 \pm 0.21$ |

The samples were collected before 9:30 a.m. to assure sample excision took place in a relaxed state, thus avoiding measurement artefacts associated with cutting under tension (Wheeler *et al*., 2013). From each of the experimental trees, one sun-exposed branch 70 - 120 cm in length or any kind of damage to avoid leaks and consequent embolism overestimation. All suspicious leakage points were sealed using a fast-drying contact adhesive (Loctite 431 with Loctite activator SF 7452, Henkel, Düsseldorf, Germany). Fruits present on the branches were removed and sealed with the same glue, as they easily detach with progressing dehydration and are hence a potential site of air-entry. Similarly, in the case of *Crataegus persimilis*, thorns were removed and sealed in the same manner to ease the handling of the branches.

### Measurements of vulnerability curves with the pneumatic method

Branch xylem vulnerability curves based on the pneumatic method (Pereira *et al*., 2016) were obtained for 44 samples from 35 experimental trees (Table 1) using the Pneumatron, a device that combines a microcontroller-activated vacuum pump and a pressure transducer to allow for automated measurements of air discharge (Pereira *et al*., 2020; Jansen *et al*., 2020).

The reservoir pressure was tracked with the Pneumatron in 0.5 sec intervals over a span of two minutes per measurement (including a pump time of approximately 2 sec in semi-automated mode and few milliseconds in automated mode). The amount of air discharged (AD) into the reservoir was calculated based on the ideal gas law (Pereira *et al*., 2016). The semi-automated mode of the Pneumatron was used to measure AD for the samples obtained from Veitshöchheim and Würzburg, whereas the additional branch samples of *T. cordata* and *T. platyphyllos* processed in Ulm were measured in automated mode. After each AD measurement, the xylem water potential was measured (see below). The maximum detectable amount of AD is associated with a change of pressure in the system by ∼50 kPa (Pereira *et al*., 2020). As the time necessary for a pressure change of 50 kPa depends on the ratio between the reservoir volume (including the cut-open conduit volume) and the volume of gas extracted from intact, embolised conduits, the choice of the optimal reservoir volume is crucial (cf. Pereira *et al*., 2020). For this study, reservoir volumes of 1.7 – 3.3 ml were selected based on the available information about the species’ embolism resistance from the xylem functional traits database (Choat *et al*., 2012), assuming that higher embolism resistance corresponds to a lower volume of gas extracted from intact, embolised conduits and hence a lower optimal reservoir volume and vice-versa. Meanwhile, however, Pereira *et al*. (2020), recommend determining the maximum reservoir volume from a completely dried branch per species. Throughout the measurements, the volume of the vacuum reservoir was kept constant for each branch.

Once the branch was fully hydrated, a clean cut was made in air at the basipetal end using a sharp razor blade to clear obstructions for air flow (Pereira *et al*., 2016; Jansen *et al*., 2020). Although this seems counterintuitive at first glance, cutting in air was done intentionally because the cut-open conduits need to be embolised when starting pneumatic measurements. The branch was then connected to the pneumatic apparatus using rigid and elastic tubing, plastic clamps and three-way stopcocks (Fig. S1, S2). The volume of the elastic tube was kept as small as possible to minimize pressure-dependent changes in reservoir volume. The elastic tubing was tightened with a plastic clamp to ensure no leakage occurred during the measurement (Bittencourt *et al*., 2018). Before each series of measurements and in case of suspicious increases in the amount of AD, the connections and the plant material were thoroughly inspected to identify and seal potential air-entry points.

Before AD measurements were taken, the branch samples were bagged in dark plastic bags to equilibrate water potential. During AD measurement, the branches were kept bagged to minimize transpiration. Between measurements, the branches were dehydrated at room temperature on a laboratory bench to induce embolism (Sperry *et al*., 1988). The branches were initially dried for intervals of about 15 – 30 min, which were subsequently increased to 1 – 4 h depending on how quickly the sample dried. To allow xylem water potential to equilibrate after each drying interval, the samples were bagged for about 30 min in the initial steps of dehydration, and for at least one hour in the later stages to account for the decrease in the leaf-stem conductance (cf. Pereira *et al*., 2016). AD measurements were made on multiple branches on the same day (see Fig. S1 – S3). The elastic tubing was always kept connected to the branch samples when switching branches to keep the reservoir volume constant throughout all measurements. AD measurements were taken until the branches were completely dehydrated and no considerable variation was observed in the amount of AD in consecutive measurements over at least 24 h, or until the maximum absolute xylem water potential measurable with the Scholander pressure chamber (10 MPa) was reached. This resulted in measurement durations of 3 – 7 days as well as in 10 – 20 AD and leaf water potential measurements per branch when following the semi-automated mode of the Pneumatron. The percentage of air discharged (PAD) was calculated as described by Pereira *et al*. (2016):

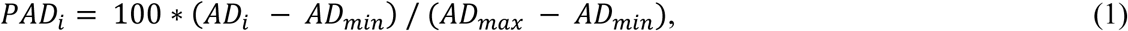

where *AD*_*i*_ is the amount of air discharged for measurement *i, AD*_*min*_ is the minimum amount of air discharged from the fully hydrated branch, and *AD*_*max*_ is the maximum amount of air discharged from the branch when completely desiccated.

### Xylem water potential measurement

For the 35 branch samples measured in semi-automated mode, the xylem pressure was measured with a Scholander pressure chamber (PMS Instruments, Corvallis, Oregon, USA) after every AD measurement. At each pressure step, two leaves were cut-off from the branch, and xylem water potential was averaged over two pressure chamber measurements. When the petiole was too small for measurement in the pressure chamber, small terminal twigs were used. The cut was immediately sealed using an instant adhesive (Loctite 431) to prevent leakage during the subsequent AD measurements. For the nine branches of *T. cordata* and *T. platyphyllos* measured using the automated mode of a Pneumatron, a stem psychrometer (ICT International, Armidale NSW Australia) was installed at a distal part of the branch to record xylem pressures for every 15 min. To obtain pressure estimates for each point in time and to reduce the impact of measurement uncertainty in the psychrometric water potentials, the psychrometer measurements for each sample were smoothed with shape-constrained additive models using monotone decreasing P-splines based on R package scam v. 1.2-8 (Pya, 2020).

### Measurements of vulnerability curves with the flow-centrifuge method

The flow-centrifuge technique (Cavitron; Cochard *et al*., 2005) was used as a reference method for comparing the agreement of xylem vulnerability curves based on hydraulic measurement methods with curves based on the Pneumatron. Flow-centrifuge measurements were performed for 49 samples from 36 trees with a Cavitron device built from a Sorval RC 5 series centrifuge with manual control of rotation speed, and using the `Cavisoft` software (`Cavisoft` version 5.2.1, University of Bordeaux, Bordeaux, France). Samples were recut several times under water to a final length of 27.5 cm to release the tension in the xylem (Torres-Ruiz *et al*., 2015). Subsequently, the non-flushed branch segments were inserted in a custom-made rotor after removing the bark at both ends. They were then spun using the principle of centrifugal force to generate a negative pressure in the xylem segment while simultaneously measuring hydraulic conductance. Flow centrifuge measurements were performed with filtered (0.2 µm) and degassed demineralized water that was enriched with 10 mM KCl and 1 mM CaCl_2_. Measurements began at xylem water potentials of around -0.8 MPa and were continued under increasingly negative xylem pressures until the percentage loss of hydraulic conductivity (PLC) reached at least 90%.

### Statistical analysis

All data handling and statistical analyses were performed in `R` version 4.0.2 (R Core Team, 2020) in the framework of the `tidyverse` (Wickham *et al*., 2019). Both the vulnerability curves based on pneumatic and flow-centrifuge measurements were described with tree-level nonlinear regression models using the logistic function by Pammenter & Van der Willigen (1998). For the flow-centrifuge method, vulnerability curves were based on the raw conductivity measured by the Cavitron (cf. Ogle *et al*., 2009):

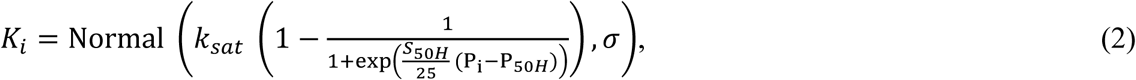

where *K*_*i*_ and *P*_i_ are the measured hydraulic conductivity and xylem pressure for observation *i*, respectively, *k*_sat_ is the hydraulic conductivity under fully saturated conditions, and *S*_50H_ is the slope at 50% loss of conductivity (*P*_50H_). For the pneumatic measurements, analogous models were constructed based on PAD for estimating *P*_50P_ (Eq. 1):

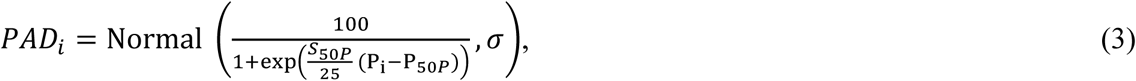

To evaluate the effect of the air-discharge time on the accuracy of the estimates of vulnerability curve parameters, separate pneumatic vulnerability curves were fit on PAD calculated for all measurement durations between the initial pressure immediately after pumping and variable final pressures in 0.5 sec intervals, from 4.5 to 115 sec. The *P*_12_ and *P*_88_ (xylem pressure at 12% and 88% loss of conductivity, respectively) were calculated from the estimated model parameters (*P*_50_ and the corresponding slope at this pressure) by rearranging the model equation (Eqs. 2 & 3), and with confidence intervals based on parametric bootstrap (*n* = 10,000). The uncertainty in VC parameters was taken into account for the calculation of species averages of VC parameters and their standard errors by using inverse-variance weighting analogous to a fixed-effects meta-analytical model (cf. Rosenberg *et al*., 2013).

We subsequently calculated a set of statistics that describe the degree of agreement of the Pneumatron parameter estimates with the flow-centrifuge based values in terms of systematic deviations, random deviations, and overall agreement. For the Pneumatron measurements from *Tilia japonica, T. cordata* and *T. platyphyllos*, these calculations were based on tree averages of the flow-centrifuge parameters when replicate measurements were performed per tree (Tab. 1).

Systematic deviations between Pneumatron and flow-centrifuge based VC parameter estimates were quantified by the mean signed deviation (MSD). The MSD only measures additive bias between the parameter estimates obtained by the pneumatic and flow-centrifuge methods (θ_P_ and θ_H_, respectively) and does not penalize scatter in the relationship.

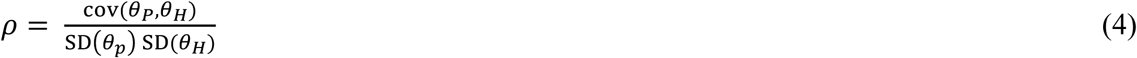

Random deviations between the estimates of both methods was evaluated by the Pearson correlation ρ between θ_P_ and θ_H_. The correlation coefficient only penalizes the degree of scatter around a hypothetical line through θ_P_ and θ_H_ and does not include information about systematic differences.

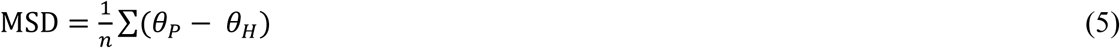

The overall agreement of the parameter estimates was evaluated by the root mean square deviation (RMSD). Unlike the aforementioned metrics, the RMSD penalizes both systematic and random deviations between θ_P_ and θ_H_.

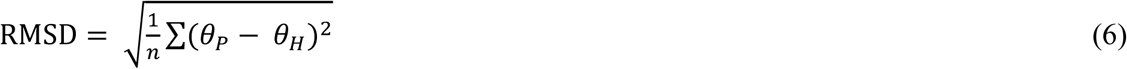

In addition, as a pragmatic measure to quantify the overall match of the flow-centrifuge and pneumatic vulnerability curves over their entire range, we calculated the L_2_ distance (cf. Cramér, 1928) between each pair of flow-centrifuge and pneumatic vulnerability curves:

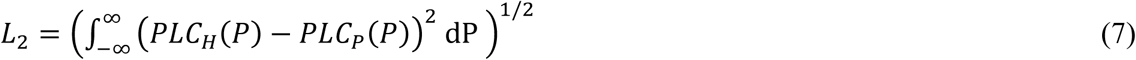

This quantity describes the degree of similarity of the two vulnerability curves and approaches zero for identical curves.

In addition to the comparison of metrics describing the agreement between estimates, we used linear mixed effects models based on R package `lme4` version 1.1-23 (Bates *et al*., 2015) to test for statistically significant differences between the Cavitron-based estimates of *P*_12_, *P*_50_, *P*_88_ and (natural log-transformed) S_50_ and their Pneumatron-based equivalents for AD intervals of 15, 30, 60, 90 and 115 sec from the initial pressure at 4 sec. Each model was fit to each parameter using restricted maximum likelihood with “method” as a fixed effect and random intercepts for species and individual trees nested in species. The flow-centrifuge was considered as baseline and contrasted with each one term for the Pneumatron estimates in five different intervals. Model assumptions were checked by residual diagnostic plots. Inference was based on Wald *t*-tests with Sattherthwaite’s approximation to the degrees of freedom using R package `lmerTest` version 3.1-2 (Kuznetsova *et al*., 2017).

### Results

### Estimated vulnerability curves

The average *P*_50H_ estimates obtained from the flow-centrifuge method covered a wide range of embolism resistance from -1.85 MPa to -6.02 MPa for *Platanus × acerifolia* and *Crataegus persimilis*, respectively. Parameters estimated with the pneumatic method largely fell into the same range (cf. Table S1). In general, there was a good agreement in overall shape between the flow-centrifuge and pneumatic vulnerability curves (VCs) for most of the species studied.

However, the estimates in *P*_12P_, *P*_50P_ and *P*_88P_ for *T. cordata* and *T. platyphyllos* based on the pneumatic method were on average at least 0.5 MPa higher than the corresponding estimates of the flow-centrifuge method when their xylem pressure was measured using a pressure chamber (Table S1). Additional measurements performed to explain this discrepancy showed that embolism resistance was largely within a comparable range when xylem pressure was determined by stem psychrometers (with the exception of *P*_88P_ for *T. cordata*; Table S1; Fig. S4). As there were reasons to assume that the differences observed were caused by inaccurate xylem water potential measurements with the pressure chamber, results for the best AD measurement intervals are based only on the observations with xylem pressure measurements based on stem psychrometers.

### Overall agreement between methods

The parameters of the vulnerability curves estimated with the two methods were generally highly correlated, with Pearson correlations above 0.54 for all parameters and over 0.74 for *P*_12_, *P*_50_ and *P*_88_ for all AD times considered in Table S2 and all 12 tree species (Table S2). In particular, this was true for the *P*_50_ estimates, where correlations exceeded 0.95 in all cases. Moreover, the *P*_50_ estimates of the two methods were very close to the 1:1 line (cf. Fig. 2b, Table S2). The Pneumatron-based estimates of the slope of the vulnerability curve, however, on average showed a negative systematic deviation that ranged from -25.5% (retransformed from log scale, 15 sec AD time) to -28.1% (115 sec AD time; cf. Fig. 1, 2d; Table S2). The deviation was notably larger in samples with lower slopes, and was not observed for the samples measured with stem psychrometers (Fig. 2). Due to the direct relationship between slope and *P*_50_, the systematically lower slope estimates translated to a higher *P*_12P_ (by up to 0.77 MPa on average at 115 sec AD time) and, in some cases, a lower *P*_88P_ (by up to -0.26 MPa on average at 15 sec AD time; Fig. 2, Table S2).

**Figure 1.**
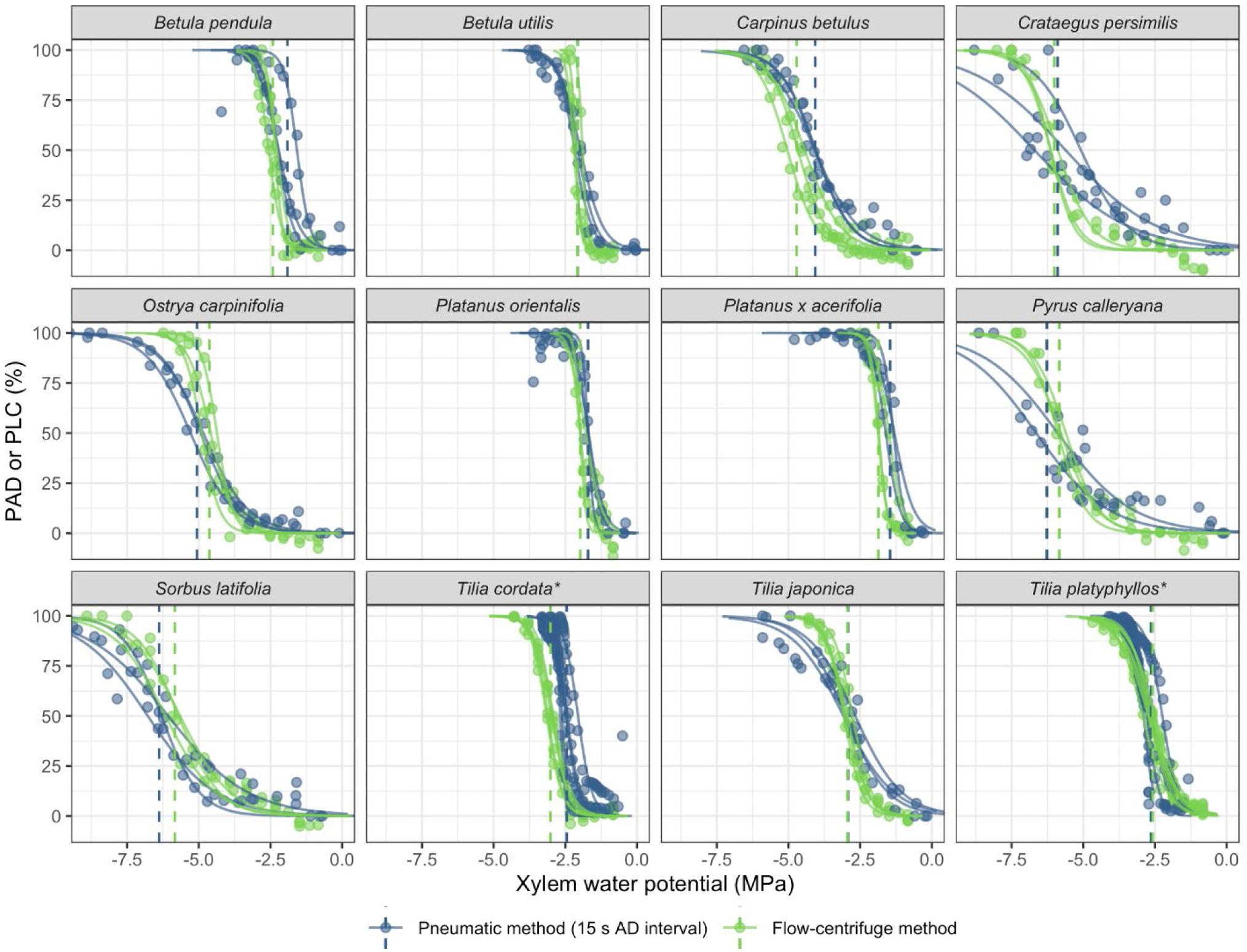
Xylem vulnerability curves obtained with the pneumatic method after a measurement duration of 15 sec (green) and the flow-centrifuge method (blue) for 12 diffuse-porous tree species. Circles: observed values (for the centrifuge data, rescaled from conductance to PLC using the estimated *k*_sat_); solid lines: predicted PLC/PAD; dashed lines: estimated *P*_50_. Asterisks (*) at the end of species names indicate xylem pressure measurements with stem psychrometers.

**Figure 2.**
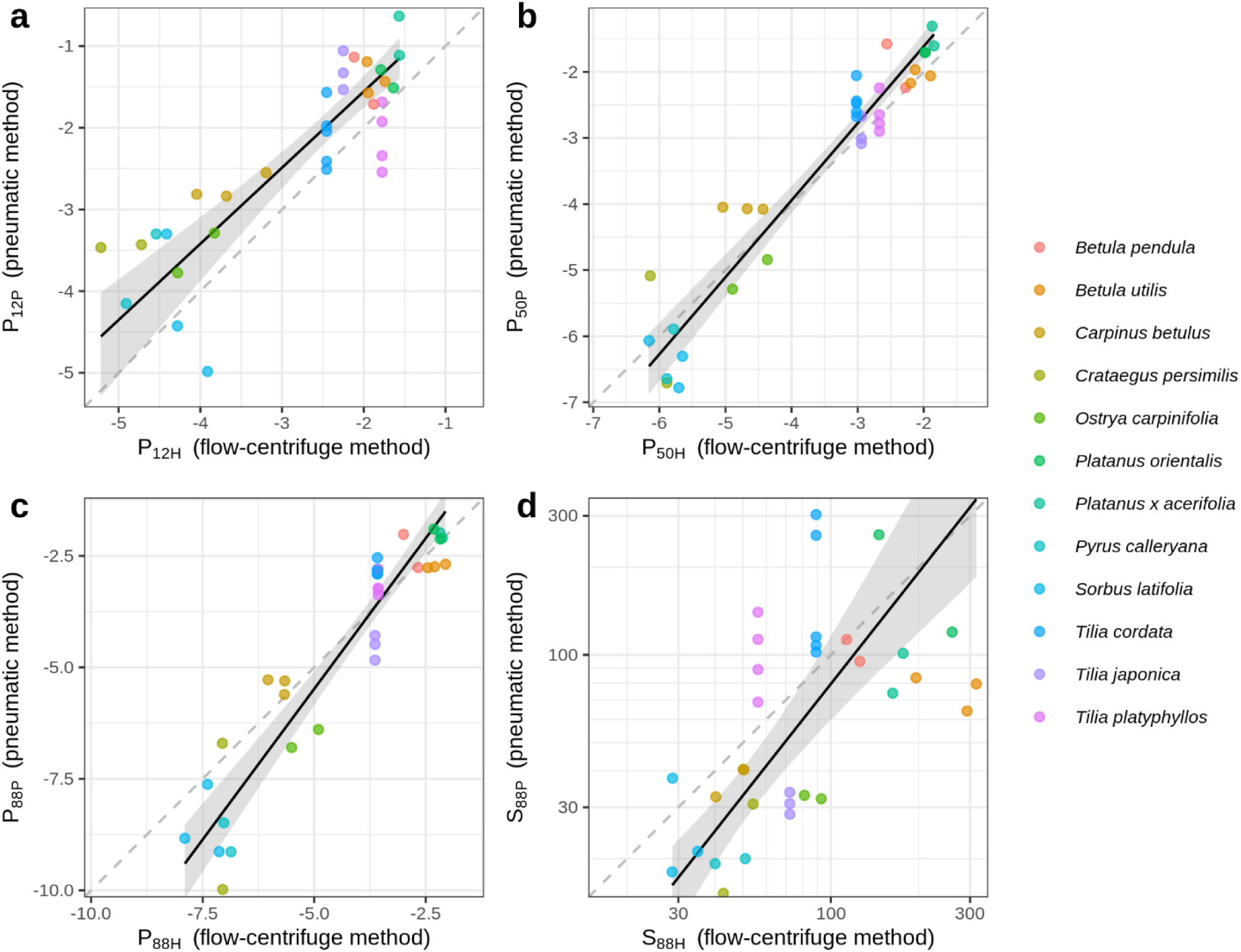
Relationship between estimates from the flow-centrifuge method (x-axis) and the pneumatic method with 15 sec air discharge interval (y-axis). a) *P*_12_, b) *P*_50_, c) *P*_88_ (xylem water potentials at 12%, 50% and 88%, respectively) and d) slope at 50% loss of conductivity (displayed on a log scale). Colours – species identity (empty circles indicate pressure chamber-based *Tilia* measurements); solid black line – standardized major axis (SMA) regression fit through all points ± 95% bootstrap confidence interval; grey dashed line: 1:1 line.

### Influence of discharge time

The pneumatic estimates of all VC parameters were sensitive to the chosen discharge time (Table 2, S2, Fig. 3, 4, 5). However, the change in agreement with air discharge time was neither consistent between parameters nor between species (Table S2, Fig. 4, 5). Consequently, the discharge interval associated with the lowest deviation between flow-centrifuge and pneumatic estimates of *P*_50_ (Fig. 3) and other parameters differed largely between species, with an increasing deviation with discharge time for some species (*Betula, Carpinus, Crataegus* and *Tilia*) and a decreasing deviation for others (*Ostrya, Platanus, Pyrus* and *Sorbus*). After accounting for random variation between species and trees, significant systematic differences between the reference value based on the flow-centrifuge method and the pneumatic method at all analysed AD times remained for all parameters, with the exception of *P*_88_ (Table 2).

**Table 2:**
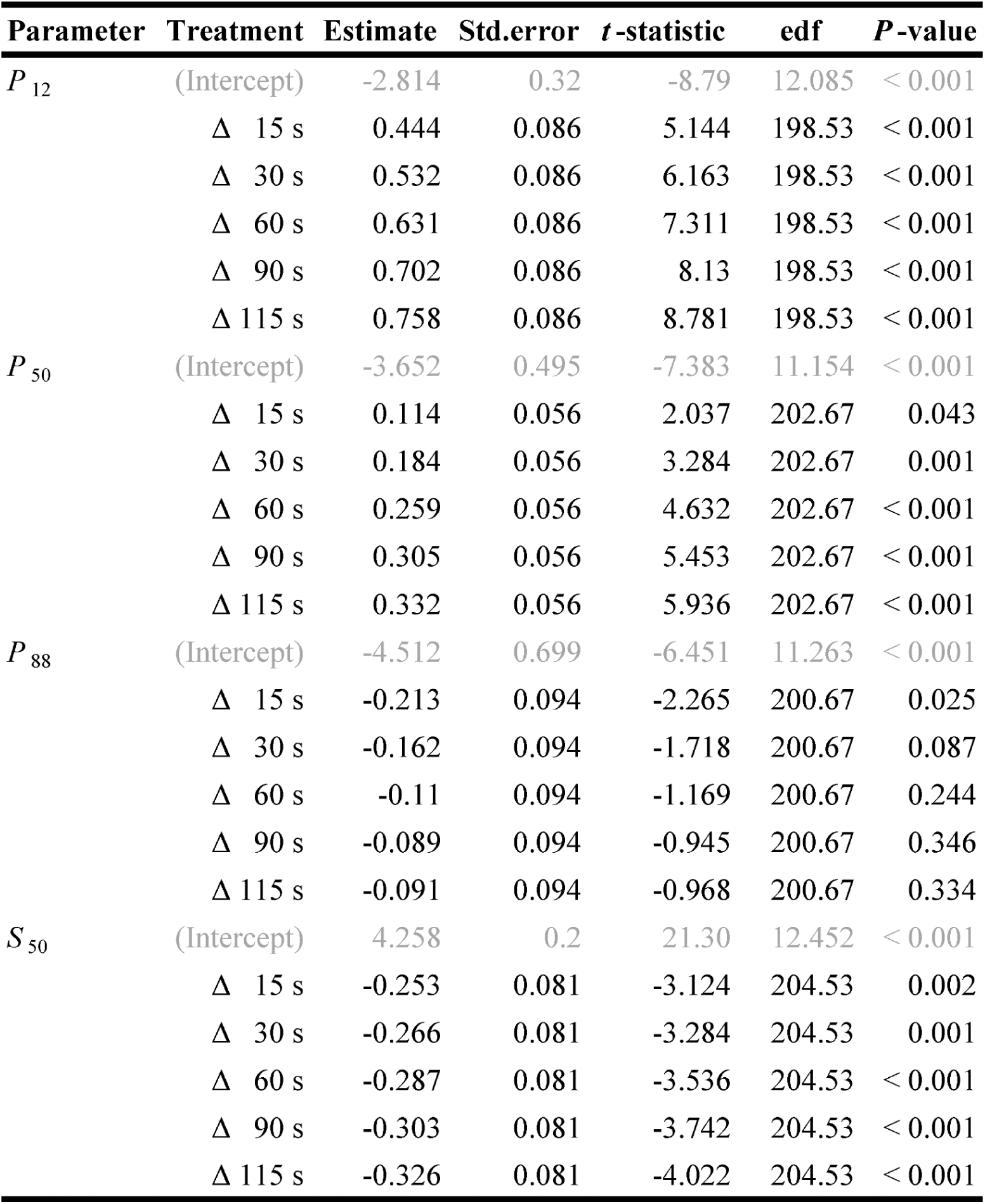
Parameter estimates of the linear mixed effect models for the xylem water potential at 12%, 50% and 88% loss of conductivity (*P*_12_, *P*_50_ and *P*_88_, respectively) and the natural log-transformed slope at 50% loss of conductivity (*S*_50_). Given are estimates for the intercept (average for flow-centrifuge based reference value) as well as the differences from the reference for different AD times, with the corresponding standard errors, *t*-statistics, approximate degrees of freedom based on the Satterthwaite approximation and corresponding *P*-values.

| Parameter | Treatment | Estimate | Std.error | $t$-statistic | edf | $P$-value |
| --- | --- | --- | --- | --- | --- | --- |
| $P_{12}$ | (Intercept) | -2.814 | 0.32 | -8.79 | 12.085 | < 0.001 |
| | $\Delta$ 15 s | 0.444 | 0.086 | 5.144 | 198.53 | < 0.001 |
| | $\Delta$ 30 s | 0.532 | 0.086 | 6.163 | 198.53 | < 0.001 |
| | $\Delta$ 60 s | 0.631 | 0.086 | 7.311 | 198.53 | < 0.001 |
| | $\Delta$ 90 s | 0.702 | 0.086 | 8.13 | 198.53 | < 0.001 |
| | $\Delta$ 115 s | 0.758 | 0.086 | 8.781 | 198.53 | < 0.001 |
| $P_{50}$ | (Intercept) | -3.652 | 0.495 | -7.383 | 11.154 | < 0.001 |
| | $\Delta$ 15 s | 0.114 | 0.056 | 2.037 | 202.67 | 0.043 |
| | $\Delta$ 30 s | 0.184 | 0.056 | 3.284 | 202.67 | 0.001 |
| | $\Delta$ 60 s | 0.259 | 0.056 | 4.632 | 202.67 | < 0.001 |
| | $\Delta$ 90 s | 0.305 | 0.056 | 5.453 | 202.67 | < 0.001 |
| | $\Delta$ 115 s | 0.332 | 0.056 | 5.936 | 202.67 | < 0.001 |
| $P_{88}$ | (Intercept) | -4.512 | 0.699 | -6.451 | 11.263 | < 0.001 |
| | $\Delta$ 15 s | -0.213 | 0.094 | -2.265 | 200.67 | 0.025 |
| | $\Delta$ 30 s | -0.162 | 0.094 | -1.718 | 200.67 | 0.087 |
| | $\Delta$ 60 s | -0.11 | 0.094 | -1.169 | 200.67 | 0.244 |
| | $\Delta$ 90 s | -0.089 | 0.094 | -0.945 | 200.67 | 0.346 |
| | $\Delta$ 115 s | -0.091 | 0.094 | -0.968 | 200.67 | 0.334 |
| $S_{50}$ | (Intercept) | 4.258 | 0.2 | 21.30 | 12.452 | < 0.001 |
| | $\Delta$ 15 s | -0.253 | 0.081 | -3.124 | 204.53 | 0.002 |
| | $\Delta$ 30 s | -0.266 | 0.081 | -3.284 | 204.53 | 0.001 |
| | $\Delta$ 60 s | -0.287 | 0.081 | -3.536 | 204.53 | < 0.001 |
| | $\Delta$ 90 s | -0.303 | 0.081 | -3.742 | 204.53 | < 0.001 |
| | $\Delta$ 115 s | -0.326 | 0.081 | -4.022 | 204.53 | < 0.001 |

**Figure 3.**
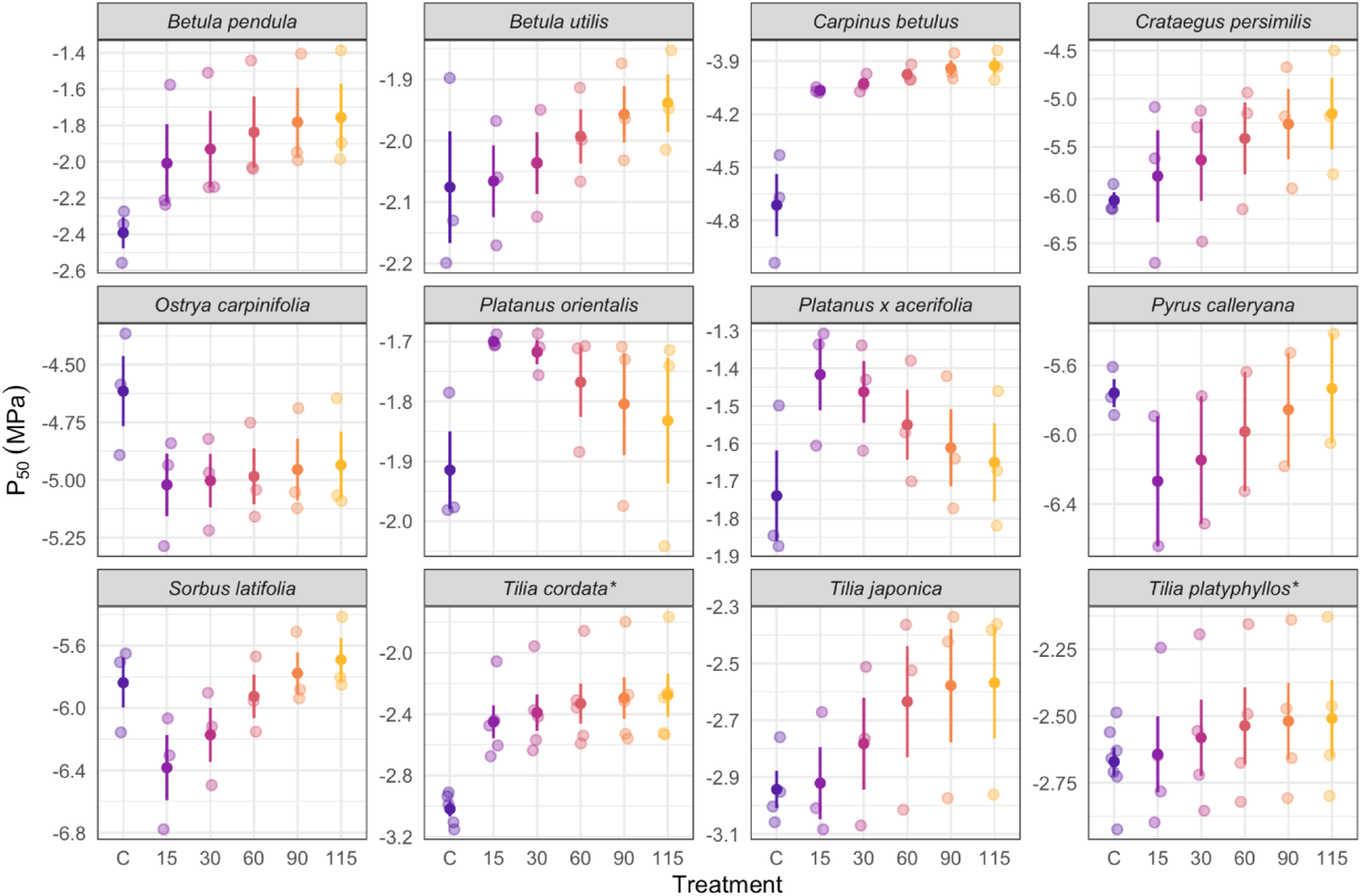
Comparison of *P*_50_ values between the flow-centrifuge and different air discharge intervals for the pneumatic method (compared on the same branch). C indicates the *P*_50_ values from the flow-centrifuge method, 15, 30, 60, 90, and 115 indicate AD measurement intervals (in seconds) evaluated from the pneumatic method. Shown are the raw estimates overlaid with their mean ± SE. Asterisks (*) at the end of species name indicate xylem pressure was determined using stem psychrometer.

**Figure 4.**
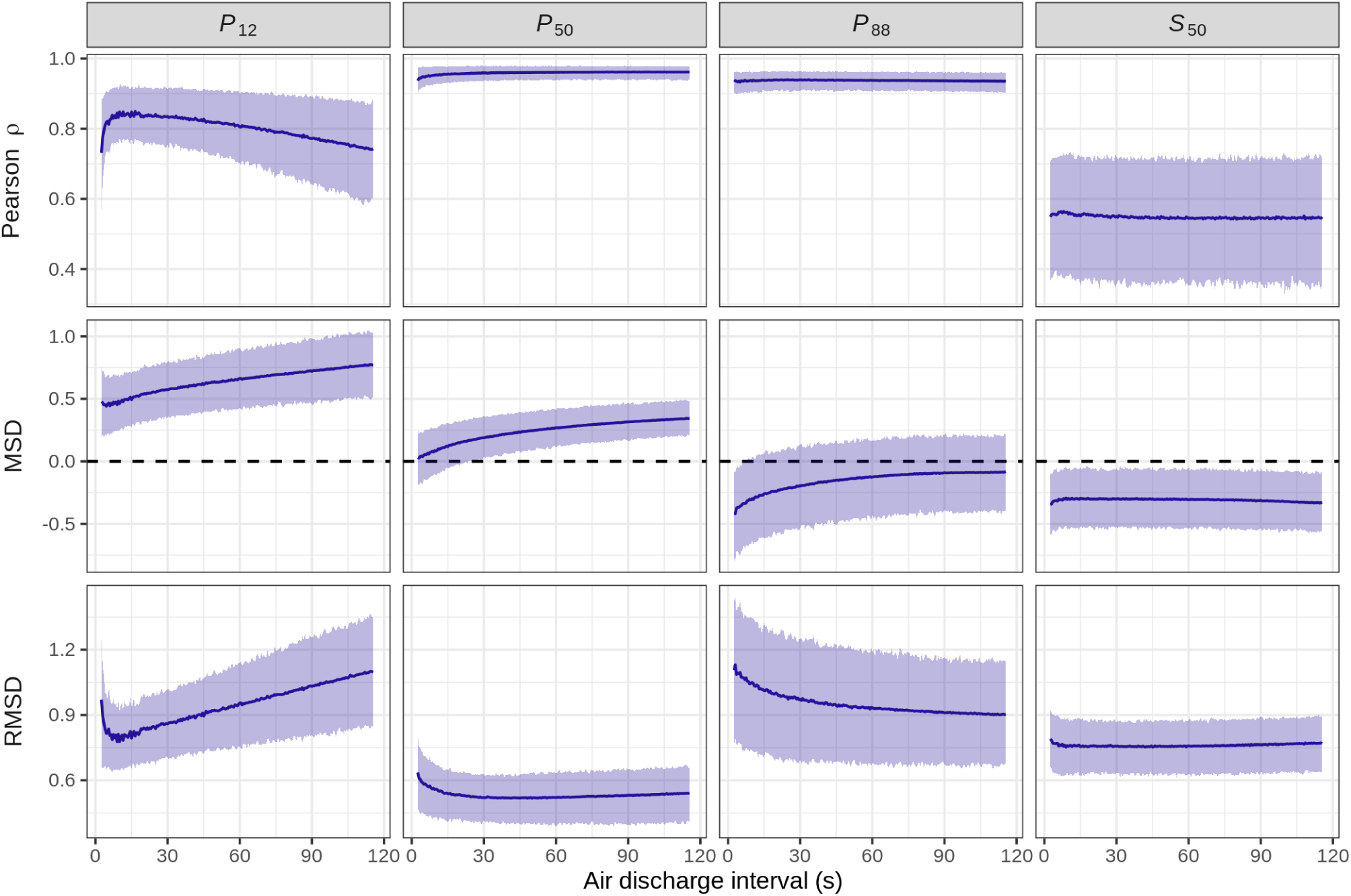
Statistics describing the agreement between the estimated vulnerability curve parameters from the pneumatic method and the flow-centrifuge method vs. the duration of air discharge measurement (estimates ± 95% bootstrap confidence intervals based on 1,000 bootstrap draws). The metrics shown are the Pearson correlation (Pearson ρ) as a measure of random deviation between the two methods (values close to one indicate a perfect linear relationship), mean signed deviation (MSD) as a measure of systematic deviations (values close to zero indicate a low bias), and root mean square deviation (RMSD) as a measure of overall agreement (low values indicate a high agreement between methods).

**Figure 5.**
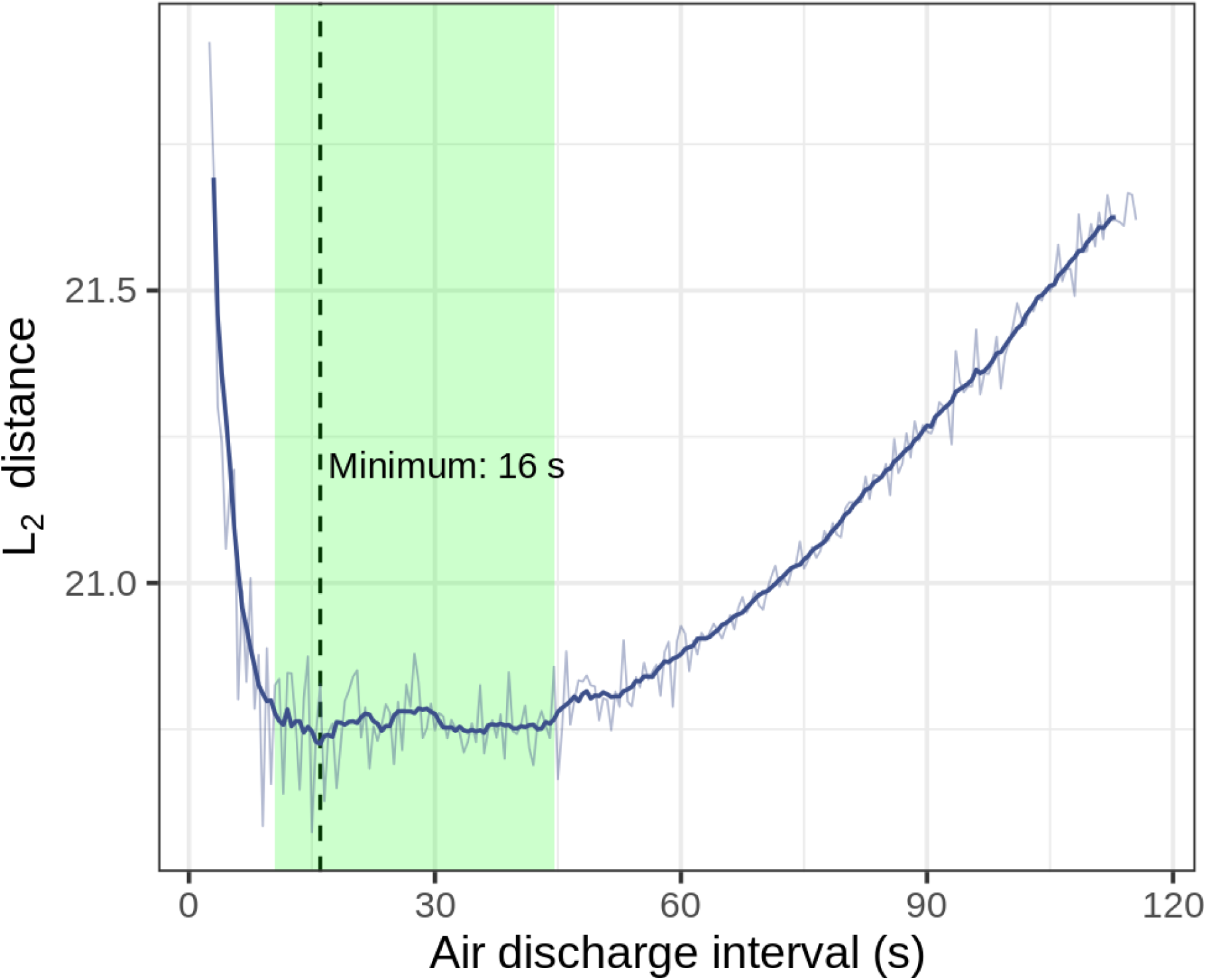
Average L_2_ distance between the pneumatic and flow-centrifuge vulnerability curves vs. air discharge time as a measure of overall accuracy (raw averages and ±5 s running average). Vertical lines indicate the minimum of the running average at 16 sec; the shaded area indicates the range with a running average differing from the minimum by less than 1% (10.5 – 44.5 sec).

The response of the pneumatic estimates of the vulnerability curve parameters to AD time was not consistent between parameters (Fig. 4). Depending on the parameter, the overall agreement with the flow-centrifuge method was either highest at low AD times (*P*_12_), remained relatively stable over a wide range of AD times (*P*_50_, *S*_50_) or increased continuously with increasing AD time (*P*_88_; cf. Fig. 4). In any case, the L_2_ distance (which describes the overall match of the curves over their entire range instead of focusing on a point estimate) was lowest at relatively low AD times with a minimum at 16 sec and a relatively broad range of equivalent values from 10.5 to 44.5 sec (Fig. 5). This indicates that the lowest degree of dissimilarity between the curves corresponded to AD times in this range, which moreover was associated with the lowest systematic differences in *P*_50_ estimates (Table 2, S2; Fig. 4).

## Discussion

In agreement with previous assessments of the pneumatic method for vulnerability curve (VC) measurements (Pereira *et al*., 2016, 2020; Zhang *et al*., 2018; Sergent *et al*., 2020), we find that the estimated water potential at 50% of discharged air volume (*P*_50P_) coincide well with the water potential at 50% loss of conductance (*P*_50H_) measured with the flow-centrifuge method for the 12 diffuse-porous temperate tree species studied, with low systematic and random deviations and a high overall agreement (MSD = +0.117 MPa, *p* = 0.956, RMSD = ±0.538 MPa, respectively for 15 sec AD time, Table S2). However, we find evidence for a high variance and systematic differences in the estimates of the slope of the VC (Fig. 1), that were on average at least 25.5% (15 sec air discharge time, cf. Table S2) lower for the pneumatic method, which also resulted in a lower agreement in the estimates of *P*_12P_ and *P*_88P_. Similar conclusions were found based on a pneumatic modelling approach (Yang *et al*., 2021), and based on experimental evidence by comparing the pneumatic and the optical method (Pereira *et al*., 2020; Guan *et al*., 2021). A mechanistic explanation for the difference in *P*_12_ and *P*_88_ values is provided by the open-xylem artefact, which suggests that embolism spreading could be enhanced by the proximity to gas under atmospheric pressure in the open-cut conduits (Guan *et al*., 2021).

Moreover, our results demonstrate that the estimates obtained with the pneumatic method are sensitive to the choice of air discharge time (cf. Pereira *et al*., 2016, 2020), to which they respond in a nonlinear and species-specific manner. For the analyzed set of species, the best overall match between the flow-centrifuge and pneumatic VCs could be achieved for air discharge times of around 15 sec (cf. Fig. 4), which experimentally confirms the ideal AD time identified by a modelling approach (Yang *et al*., 2021).

### Agreement between flow-centrifuge and pneumatic vulnerability curves

The gradual loss of conductance and the increasing amount of gas extracted, which are quantified by the PLC and PAD, are necessarily positively associated as they share a common cause, namely embolism spreading under progressing dehydration. However, PAD quantifies the air volume inside embolised vessels, while PLC measures their contribution to the conductance of the branch. Thus, both arise from vessel properties that scale with different powers of their length and diameter (Pereira *et al*., 2016). While the volume of a vessel scales with the second power of diameter and is proportional to its length, its flow resistance (i.e. the inverse of its conductance) can be approximated by the sum of the resistance posed by its lumen and the resistance posed by the transition through pit membranes (Sperry *et al*., 2005; Wheeler *et al*., 2005). According to the Hagen-Poseuille-equation, the flow resistance of the lumen scales with the inverse of the fourth power of its diameter and is proportional to its length. The flow resistance posed by the transition through pit membranes can be assumed to scale with the inverse of expected vessel length (Sperry *et al*., 2005). Due to these different scaling relationships, the association between PAD and PLC does not necessarily have to be linear. Observed systematic differences between the parameters of curves obtained with different methods (cf. Table 2) might therefore not be surprising and deserve further testing.

It should also be noted that an additional downward bias in the slope might be introduced by treating the unknown true minimum and maximum amount of air discharged (*AD*_min_ and *AD*_max_, cf. Eq. 1) as fixed quantities measured without error. For hydraulic vulnerability curves, problems induced by treating maximum conductivity as fixed can be circumvented by treating the saturated hydraulic conductivity as a model parameter (*k*_sat_ in Eq. 2; Ogle *et al*., 2009; Duursma & Choat, 2017). Implementing a similar solution for the pneumatic method is not straightforward due to the common identifiability issues in sigmoidal models where the lower and upper bound are both treated as model parameters.

Given the direct mathematical relation between the model parameters, a bias in slope can be expected to result in a bias in *P*_12_ and/or *P*_88_, which may be problematic in process-based vegetation modelling that use these VC parameters to describe drought resistance. Our results indicate that while the pneumatic method produces reliable estimates of the *P*_50_ for diffuse-porous tree species, the curves obtained might not always be interchangeable with curves constructed with the flow-centrifuge method.

### Effect of the air discharge interval on measurement accuracy

The observed sensitivity of the pneumatic method to the AD interval provides experimental evidence that is in line with predictions by the Unit Pipe Pneumatic model (Yang *et al*., 2021). To understand the relationship between the AD interval and the amount of air discharged in that interval, it is necessary to focus on the underlying assumptions about the gas flow between sample and reservoir. One central assumption of the pneumatic method is that the amount of air discharged into the vacuum reservoir over a given time span is a function of the amount of air *N* inside embolised conduits at different points during the dehydration process (Jansen *et al*., 2020; Yang *et al*.,2021). This assumption is necessary to be able to use PAD measurements to infer the degree of embolization. Further, if the PAD calculated over different AD times *t* is to result in identical vulnerability curves, the change in AD with time must have the same shape at different dehydration steps and only differ by a multiplicative constant proportional to the amount of air *N* in the xylem, i.e. AD(*t, N*_1_) / *N*_1_ = AD(*t, N*_2_) / *N*_2_ must hold. This assumption is likely only approximately met under typical measurement conditions. During the drying process, the xylem undergoes substantial changes that may affect the shape of AD(*t, N*), such as an increase in gas conductivity with progressing embolism. Moreover, it is possible that embolised conduits that were not disconnected from to the cut surface at earlier drying steps subsequently become connected to the network of open vessels connecting to the vacuum reservoir (Pereira et al. 2016). In such cases, the amount of air in these spaces would not be included in the earlier estimates. For these reasons, the relationship between the AD measured in different desiccation steps after a certain discharge interval and the total amount of air within the xylem at those points in time is likely empirical. This may explain the pronounced species differences in the response to AD time (Table S2, 4, Fig. 3), and contribute to the previously reported differences in accuracy for species with different types of wood anatomy (e.g. Zhang *et al*., 2018).

While the effect of AD time depended on species identity, and was different for different parameters, Fig. 4 indicates that overall mismatch between flow-centrifuge and pneumatic VCs could be minimized by choosing low AD times of ca.16 sec. In our setting, this might be a consequence of small amounts of air-entry into the xylem where the branch is damaged, which– unlike the other discussed factors influencing AD – will have an effect that accumulates with discharge time. Thus, the net gains in accuracy due to integrating over a larger time interval are overcompensated by the increasing contribution of potential leakage to the total AD. Interestingly, the higher agreement with hydraulic reference values at lower AD times contrasts with the higher accuracy for longer AD times reported in earlier works using the manual pneumatic method (cf. Fig. S3 in Pereira et al. 2016). Most likely, this difference results from the higher temporal resolution and more accurate AD time measurement enabled by the automated Pneumatron device (Pereira et al. 2020). When using a Pneumatron, the choice of short AD times therefore is a pragmatic way to improve the accuracy of the pneumatic method.

### Species-specific drying behaviour

As noted previously (Zhang *et al*., 2018; Sergent *et al*., 2020), the degree of similarity between VCs measured with the pneumatic method and hydraulic reference methods differs between species. In this study, we observed a mismatch for *T. cordata* and *T. platyphyllos* when xylem pressure was expressed based on leaf water potentials measured with a pressure chamber. However, the difference largely disappeared when a stem psychrometer was used to determine xylem pressure (Fig. 1, S4).

Possible explanations for the observed differences is the presence of abundant mucilage in the xylem tissues of *Tilia* species (Franz & Kram, 1985; Pigott *et al*., 2012), which has been reported to affect xylem pressures measured with the pressure chamber technique (Zimmermann *et al*., 2002). Furthermore, it could be speculated that the truncated shape of the pressure-chamber based Pneumatron VCs for the *Tilia* species (especially *T. cordata*, cf. Fig. S4), and the fact they never reached water potentials substantially more negative than -3 MPa, indicates that the measurements were terminated before the stems of the specimens were fully dehydrated. This results in underestimated values for *AD*_max_, thus shifting the curve towards less negative water potentials. As detailed before, the desiccation was continued until a near constancy in the measured *AD* values over at least 24 h was reached. This criterion to finish *AD* measurements may not be ideal, as the drying process may slow down notably after full stomatal closure, which especially for isohydric species may happen in a relatively well-hydrated state. Notably, while *Tilia* species have often been considered to be relatively anisohydric (cf. Leuzinger *et al*., 2005; Galiano *et al*., 2017; Kiorapostolou *et al*., 2018; but see Niinemets *et al*., 1999), recent work by Leuschner *et al*. (2019) indicates that at least *T. cordata* has a fairly stringent, more isohydric stomatal control mechanism. Due to the anticipated hydraulic segmentation between leaf petioles and stem xylem in drought-avoiding and more isohydric species (Hartmann *et al*., 2021), this may have contributed to the mismatch between flow-centrifuge and pneumatic VCs. It is well documented that certain species rely on early drought-induced leaf shedding (Wolfe *et al*., 2016; Hochberg *et al*., 2017), most likely caused by a pronounced hydraulic segmentation (cf. Pivovaroff *et al*., 2016; Zhu *et al*., 2016; Klepsch *et al*., 2018). As these processes decouple leaf and branch water potentials, measuring leaf water potential with a Scholander pressure chamber may result in extreme water potential readings that do not reflect the actual status in the xylem. Conversely, the slowdown of dehydration induced by leaf shedding may result in prematurely terminated measurements when measuring branch water potentials with e.g. a stem psychrometer, or when determining the end of the dehydration process based on the state of the leaves. As a cautionary example, the leaves of *C. persimilis* were almost fully dehydrated on the third day of measurements, while the branch water potential values continued to decline for seven days. Similar behaviour was reported by Wolfe *et al*. (2016) for the tropical species *Genipa americana*.

Due to these species-specific differences in drying behaviour, the stability of *AD*_max_ can have a strong influence on the shape of the vulnerability curves because all pneumatic *PAD* values are normalized against *AD*_min_ and *AD*_max_. It may therefore be advantageous to continue the drying process until the constancy of *AD*_max_ has been confirmed based on several measurements to avoid bias resulting from underestimating *AD*_max_. An important corollary of the observed problems associated with water potential measurements is that the same kind of bias in water potential may also affect hydraulic VC measurements in other methods that rely on bench dehydration.

### Implications for future vulnerability curve method comparisons

An important limitation that affects all evaluation studies of VC methods – as well as most other measurement methods in biology – is the lack of true reference values for VC parameters. While in this study, we used measurements with the flow-centrifuge method as a reference, there are many indications that this method may be affected by measurement artefacts that arise during sample excision (Wheeler *et al*., 2013) and preparation (Torres-Ruiz *et al*., 2015) or as a result from vessel lengths exceeding the sample dimensions (Choat *et al*., 2010; Martin-StPaul *et al*., 2014; Torres-Ruiz *et al*., 2014). A mismatch between curves obtained with the pneumatic method and the flow-centrifuge method may thus in part also be attributed to the imperfections of the latter. While there is hardly a way to overcome the limitation of imperfect reference values, we argue that methodological comparisons of VC methods may benefit from adopting a more principled approach of quantifying the agreement between different methods in terms of different components of accuracy (cf. Fuchs *et al*., 2017; Flo *et al*., 2019). To our knowledge, none of the previously published methodological comparisons (see e.g. Li *et al*., 2008; Choat *et al*., 2010; Hacke *et al*., 2015; Brodribb *et al*., 2017; López *et al*., 2019; Venturas *et al*., 2019; Chen *et al*., 2021; Pratt *et al*., 2020; Sergent *et al*., 2020; Zhao *et al*., 2020) formally differentiate between systematic and random differences in parameter estimates, and none provide metrics that quantify the similarity over the entire curves. While our choice of the L_2_-distance as a measure of overall agreement between curves is relatively arbitrary and there are many equally appropriate distance metrics, it is most definitely an improvement compared to the common practice of comparing methods by the Pearson correlation between parameter estimates, as the latter only penalizes deviations from a bivariate linear relationship while being insensitive to systematic deviations. We hope that our framework can serve as a starting point for more formal VC method comparisons based on rigorous metrological principles and theory.

## Conclusions

Our data indicate a high degree of agreement between the *P*_50P_ estimated with the pneumatic method and the *P*_50H_ estimated with the flow-centrifuge method for the analysed diffuse-porous temperate tree species, especially when using short air discharge times of around 15 sec. The relatively low effort required to construct a curve with this method and its high degree of automation when using a Pneumatron device in conjunction with a stem psychrometer allow for a high throughput. The method is therefore attractive in descriptive or predictive contexts where the main purpose is to generate a good proxy for plant drought resistance. However, the observed systematic deviation in slope estimates as well as potential artefacts associated with xylem water potential determination and species-specific drying behaviour deserve further attention.

## Acknowledgements

We thank Klaus Körber and Andreas Lösch from the Bavarian state institute for viticulture and horticulture (Bayerische Landesanstalt für Wein-und Gartenbau, LWG) for granting access to their research facility at Stutel and Xinyi Guan for her support with the automated pneumatic measurements. P.R.L.B. acknowledges Royal Society’s Newton International for its Fellowship (NF170370). S.J. acknowledges funding from the German Research Foundation (Deutsche Forschungsgemeinschaft, DFG, project nr. 410768178).

## Author contributions

B.S. and R.M.L. designed the study, S.S.P. performed the semi-automated pneumatic measurements, technically supported by P.B. and L.P., L.P. the automated pneumatic measurements and E.I. the hydraulic measurements. S.S.P. and R.M.L. analysed the data. S.S.P, R.M.L. and B.S. wrote the first manuscript, which was intensively discussed and revised by all authors.

