## Supplementary tables and figures for "Accuracy of the pneumatic method for estimating xylem vulnerability to embolism in temperate diffuse-porous tree species"

**Table S1:** Comparison of the estimated xylem water potential at 12%, 50% and 88% loss of conductivity / percent air discharged ($P_{12}$, $P_{50}$ and $P_{88}$, respectively) and slope of the vulnerability curve ($S_{50}$) for the 12 studied diffuse-porous tree species for the pneumatic (15 sec AD interval) and flow-centrifuge method. Given are the means and propagated standard errors including the inferential uncertainty of the PLC measurements. Asterisks (*) indicate xylem pressures measured with stem psychrometers; hashes (#) indicate individuals from the same species measured with the pressure bomb.

| | $P_{12}$ (MPa) | | $P_{50}$ (MPa) | | $P_{88}$ (MPa) | | $S_{50}$ (-% $MPa^{-1}$) | |
|---|---|---|---|---|---|---|---|---|
| **Species** | **Cavitron** | **Pneu. (15 s)** | **Cavitron** | **Pneu. (15 s)** | **Cavitron** | **Pneu. (15 s)** | **Cavitron** | **Pneu. (15 s)** |
| *Betula pendula* | -2.07 ± 0.03 | -1.36 ± 0.05 | -2.38 ± 0.01 | -1.85 ± 0.02 | -2.69 ± 0.03 | -2.48 ± 0.05 | 134.1 ± 10.5 | 77.70 ± 5.53 |
| *Betula utilis* | -1.92 ± 0.01 | -1.37 ± 0.05 | -2.09 ± 0.00 | -2.05 ± 0.02 | -2.24 ± 0.01 | -2.73 ± 0.05 | 267.0 ± 11.9 | 70.74 ± 4.52 |
| *Carpinus betulus* | -3.74 ± 0.06 | -2.65 ± 0.08 | -4.75 ± 0.03 | -4.06 ± 0.03 | -5.76 ± 0.05 | -5.48 ± 0.08 | 48.03 ± 2.12 | 34.24 ± 1.63 |
| *Crataegus persimilis* | -4.96 ± 0.17 | -3.39 ± 0.21 | -6.02 ± 0.06 | -5.62 ± 0.09 | -7.05 ± 0.16 | -7.56 ± 0.23 | 46.90 ± 5.44 | 17.23 ± 1.50 |
| *Ostrya carpinifolia* | -4.09 ± 0.04 | -3.57 ± 0.06 | -4.72 ± 0.02 | -4.99 ± 0.03 | -5.32 ± 0.04 | -6.44 ± 0.07 | 75.68 ± 4.00 | 34.14 ± 1.39 |
| *Platanus acerifolia* | -1.55 ± 0.02 | -1.00 ± 0.05 | -1.85 ± 0.01 | -1.42 ± 0.02 | -2.14 ± 0.02 | -1.97 ± 0.04 | 154.7 ± 8.00 | 95.95 ± 8.07 |
| *Platanus orientalis* | -1.73 ± 0.04 | -1.50 ± 0.04 | -1.97 ± 0.02 | -1.70 ± 0.02 | -2.25 ± 0.04 | -1.91 ± 0.04 | 135.9 ± 16.0 | 131.2 ± 17.6 |
| *Pyrus calleryana* | -4.73 ± 0.10 | -3.99 ± 0.28 | -5.79 ± 0.04 | -6.37 ± 0.14 | -6.82 ± 0.09 | -8.97 ± 0.41 | 45.94 ± 3.42 | 19.76 ± 2.28 |
| *Sorbus latifolia* | -3.94 ± 0.18 | -4.34 ± 0.15 | -5.69 ± 0.07 | -6.33 ± 0.06 | -7.39 ± 0.14 | -8.19 ± 0.14 | 28.99 ± 2.17 | 20.34 ± 1.27 |
| *Tilia cordata** | -2.45 ± 0.02 | -2.26 ± 0.00 | -3.02 ± 0.01 | -2.55 ± 0.00 | -3.60 ± 0.02 | -2.84 ± 0.00 | 84.42 ± 2.68 | 114.7 ± 1.03 |
| *Tilia cordata#* | -2.45 ± 0.04 | -1.47 ± 0.07 | -3.15 ± 0.02 | -2.22 ± 0.03 | -3.85 ± 0.04 | -3.03 ± 0.06 | 55.97 ± 2.84 | 61.48 ± 4.31 |
| *Tilia japonica* | -2.35 ± 0.02 | -1.35 ± 0.15 | -3.00 ± 0.01 | -2.93 ± 0.05 | -3.64 ± 0.02 | -4.53 ± 0.14 | 74.41 ± 1.69 | 30.79 ± 2.41 |
| *Tilia platyphyllos** | -1.82 ± 0.03 | -2.21 ± 0.01 | -2.68 ± 0.01 | -2.71 ± 0.00 | -3.54 ± 0.02 | -3.20 ± 0.01 | 55.52 ± 1.51 | 83.75 ± 0.88 |
| *Tilia platyphyllos#* | -2.95 ± 0.02 | -1.44 ± 0.05 | -3.43 ± 0.01 | -1.92 ± 0.02 | -3.92 ± 0.02 | -2.42 ± 0.05 | 82.46 ± 3.17 | 90.53 ± 8.14 |

**Table S2:** Summary of a set of metrics describing the agreement between the two analyzed methods for the estimated parameters for the xylem water potential at 12%, 50% and 88% loss of conductivity ($P_{12}$, $P_{50}$ and $P_{88}$, respectively), and the slope at 50% loss of conductivity ($S_{50}$). Given are the comparisons between the flow-centrifuge method and the pneumatic method at 15, 30, 60, 90 and 115 sec air discharge intervals. The values indicating the highest agreement are highlighted in bold.

| Parameter | AD time | Pearson ρ | MSD | RMSD |
|---|---|---|---|---|
| $P_{12}$ | 15 s | **0.848(0.778−0.919)** | **0.493( 0.274− 0.695)** | **0.793(0.654−0.932)** |
| | 30 s | 0.835(0.757−0.913) | 0.574( 0.352− 0.801) | 0.859(0.704−1.018) |
| | 60 s | 0.805(0.700−0.900) | 0.662( 0.430− 0.902) | 0.955(0.753−1.138) |
| | 90 s | 0.772(0.644−0.891) | 0.720( 0.455− 0.982) | 1.032(0.807−1.256) |
| | 115 s | 0.741(0.605−0.881) | 0.771( 0.507− 1.029) | 1.099(0.852−1.342) |
| $P_{50}$ | 15 s | 0.956(0.933−0.978) | **0.117(-0.058− 0.307)** | 0.538(0.408−0.646) |
| | 30 s | 0.959(0.937−0.978) | 0.189( 0.038− 0.362) | 0.520(0.419−0.627) |
| | 60 s | 0.961(0.938−0.979) | 0.269( 0.110− 0.410) | **0.520(0.386−0.632)** |
| | 90 s | 0.961(0.938−0.978) | 0.315( 0.162− 0.465) | 0.531(0.399−0.651) |
| | 115 s | **0.962(0.939−0.978)** | 0.343( 0.208− 0.484) | 0.540(0.411−0.659) |
| $P_{88}$ | 15 s | 0.938(0.909−0.963) | -0.259(-0.616− 0.060) | 1.011(0.701−1.324) |
| | 30 s | **0.939(0.908−0.962)** | -0.196(-0.534− 0.137) | 0.972(0.697−1.251) |
| | 60 s | 0.938(0.902−0.961) | -0.125(-0.421− 0.170) | 0.933(0.681−1.185) |
| | 90 s | 0.936(0.906−0.961) | -0.090(-0.389− 0.208) | 0.909(0.672−1.165) |
| | 115 s | 0.935(0.903−0.960) | **-0.085(-0.400− 0.223)** | **0.902(0.673−1.147)** |
| $\log(S_{50})$ | 15 s | **0.551(0.351−0.716)** | **-0.295(-0.511−-0.044)** | **0.754(0.624−0.879)** |
| | 30 s | 0.550(0.357−0.721) | -0.300(-0.525−-0.054) | 0.756(0.635−0.863) |
| | 60 s | 0.544(0.353−0.707) | -0.306(-0.539−-0.048) | 0.758(0.630−0.878) |
| | 90 s | 0.544(0.358−0.718) | -0.312(-0.527−-0.058) | 0.763(0.636−0.893) |
| | 115 s | 0.545(0.343−0.723) | -0.330(-0.556−-0.090) | 0.772(0.635−0.892) |

**Figures:**

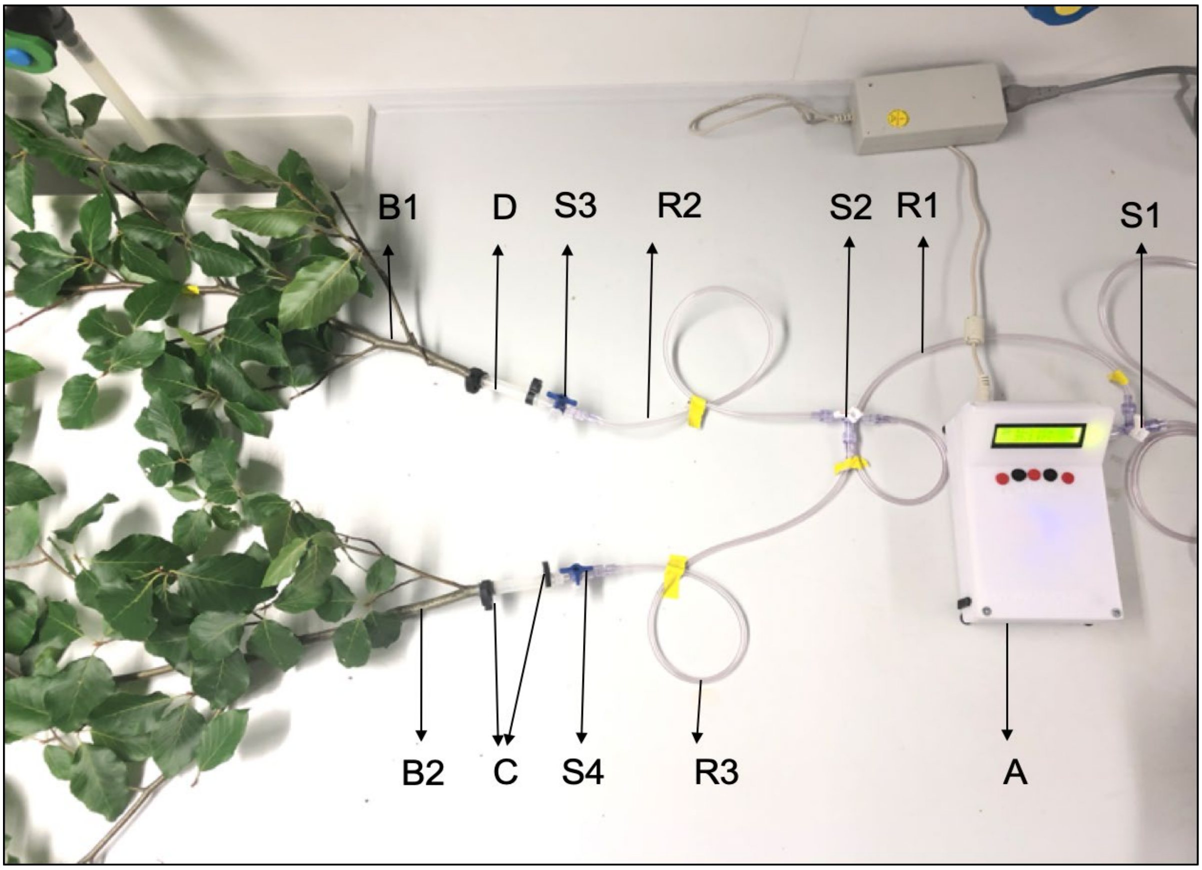


**Figure S1:** Documentation of the applied set-up. A – Pneumatron: device for air discharge measurements; B1 and B2: – branch 1 and branch 2; C – Plastic clamps to tighten the connection of elastic tubing and branch to avoid leakage; D – Elastic tube to connect the branch to the vacuum reservoir; R1, R2, R3 – Vacuum reservoir; S1, S2, S3 and S4 – three-way stopcocks for switching the connection between the pneumatic apparatus and the two branches.

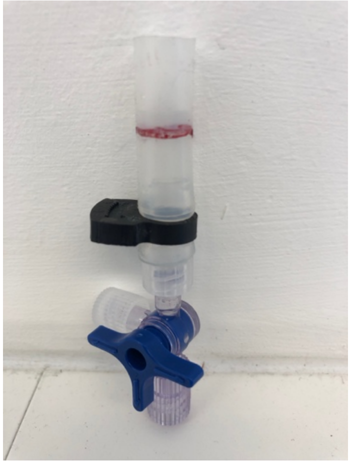

**Figure S2:** Documentation of the stopcock, plastic clamps, and elastic tubing used for connecting the pneumatic apparatus with rigid tubing to the branch. The red color marks the point where the branch was inserted into the tubing. To calculate the volume of elastic tubing, the empty weight of stopcock, plastic clamps, and elastic tubing was measured and filled up to the red mark with water after measurement and weighed again. The weight of the water was later converted into the corresponding volume at the temperature at the time of measurement.

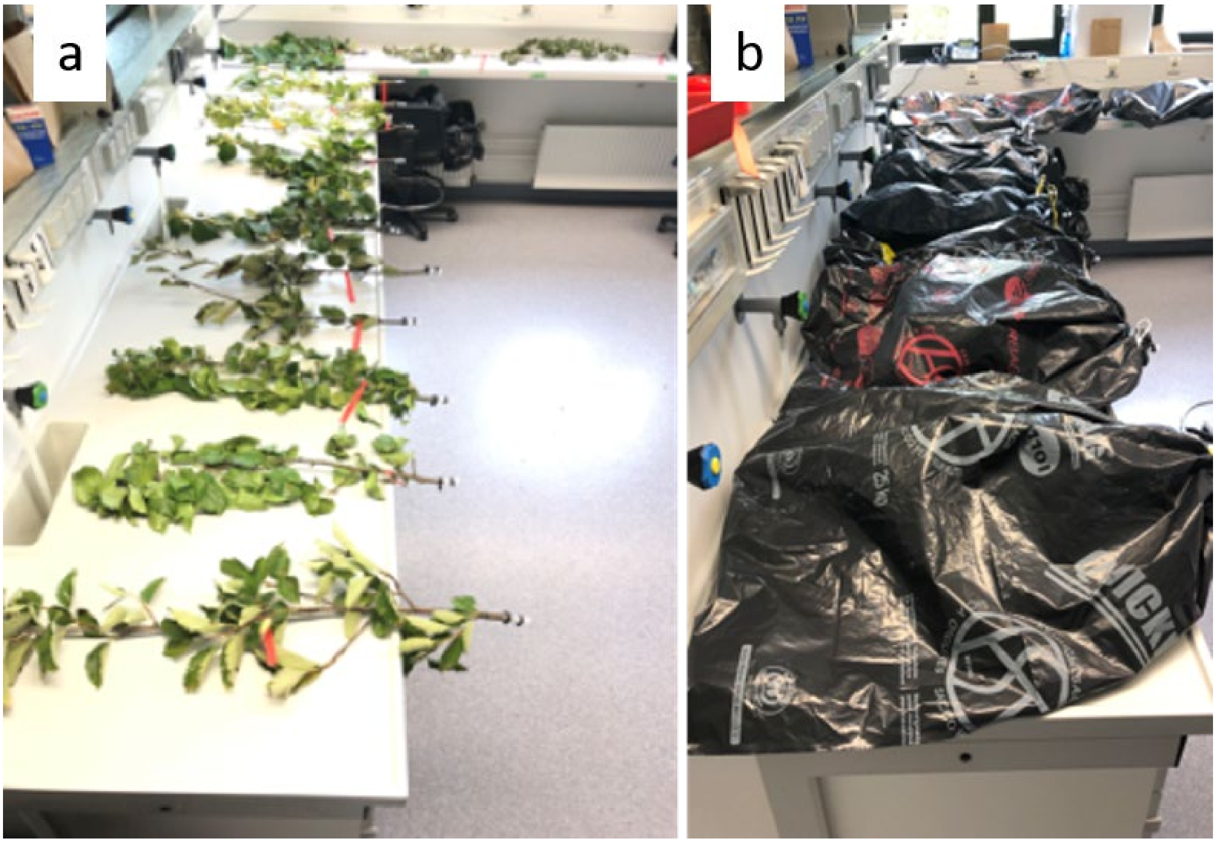


**Figure S3:** Storage of branch samples during the measurement. a) Samples being dried on the bench to induce embolism. Drying intervals between air discharge measurements varied from about 15 min at the initial stages to 6-8 h at higher desiccation levels. All of these branches were measured simultaneously, which highlights the advantage of the high sample throughput of the pneumatic method. b) Branch samples kept for equilibration by covering them in black plastic bags before measurement. Initially, the branches were kept for 30 min for equilibration before each measurement, which was increased to at least one hour at the later stages.

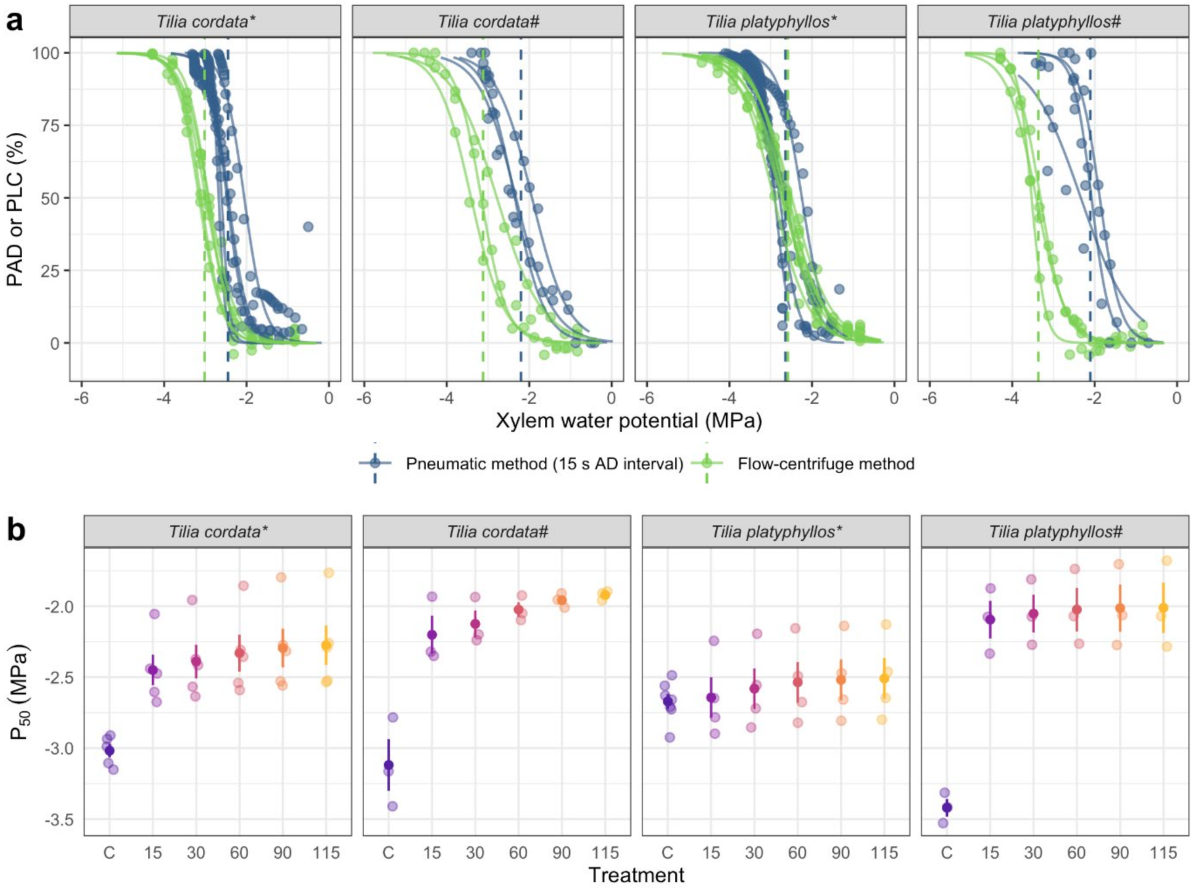


**Figure S4:** Comparison of VC and $P_{50}$ estimates measured using stem psychrometer and pressure chamber a) Xylem vulnerability curves obtained with the pneumatic method for 15 sec air discharge interval (green) and curves from the flow-centrifuge method (blue) for two *Tilia* species. Circles: observed values (for the centrifuge data, rescaled from conductance to PLC using the estimated $k_{sat}$); solid lines: predicted PLC/PAD; dashed lines: estimated $P_{50}$. b) Comparison of $P_{50}$ values between the flow-centrifuge and different air discharge intervals for the pneumatic method (compared on the same branch). C indicates the $P_{50}$ values from the flow-centrifuge method, 15, 30, 60, 90, and 115 indicate AD measurement intervals evaluated from the pneumatic method. Shown are the raw estimates overlaid with their mean ± SE. Asterisks (*) indicate xylem pressures measured with stem psychrometers; hashes (#) indicate individuals from the same species measured with the pressure bomb.
